# Glutaredoxins rapidly reduce glutathione hydroper- and polysulfides

**DOI:** 10.64898/2026.07.01.735490

**Authors:** Philipp Reinert, Seiryo Ogata, Laura Leiskau, Sernur Sena Yildiz, Takaaki Akaike, Uladzimir Barayeu, Marcel Deponte

**Affiliations:** Department of Chemistry, Comparative Biochemistry, RPTU Kaiserslautern, D-67663 Kaiserslautern, Germany; Max-Planck-Institute for Polymer Research, D-55128 Mainz, Germany; Department of Redox Molecular Medicine, Tohoku University Graduate School of Medicine, Sendai 980-8575, Japan

**Keywords:** Glutaredoxin, polysulfide, hydropersulfide, hydrogen sulfide, hydrogen polysulfides

## Abstract

Hydropersulfides have gained attention in cell biology as excellent nucleophiles and membrane-protective radical scavengers. They form perthiyl radicals, which terminate radical chain reactions through self-recombination, leading to the formation of polysulfides. It is currently unknown how polysulfides are subsequently reduced again in non-enzymatic or enzymatic metabolic pathways. Here we used stopped-flow kinetic measurements in combination with mass spectrometry to show that the model class I glutaredoxin from the malaria parasite *Plasmodium falciparum* (PfGrx) rapidly reduces the polysulfides glutathione trisulfide (GS_3_G) and glutathione tetrasulfide (GS_4_G), yielding the glutathionylated enzyme and the corresponding glutathione hydropersulfide GSSH and hydrotrisulfide GS_3_H. The second-order rate constants of these enzymatic reductions ≥10^7^ M^−1^s^−1^ are even slightly higher than for glutathione disulfide (GSSG). In contrast, PfGrx was inactive or only moderately active using cystine or cysteine trisulfide as oxidants. GSSH and GS_3_H are further reduced by PfGrx with second-order rate constants on the order of 10^6^–10^7^ M^−1^s^−1^, yielding the glutathionylated enzyme as well as hydrogen sulfide (H_2_S) and hydrogen disulfide (H_2_S_2_), respectively. Thus, glutaredoxins specifically recognize the glutathione moiety of glutathione (hydro)polysulfides and glutathione hydropersulfide. Due to the rapid reduction of glutathionylated glutaredoxins by reduced glutathione (GSH), glutathione (hydro)per/polysulfides are efficiently converted to GSSG and H_2_S or the corresponding hydrogen polysulfides. As a consequence, the steady-state concentration of glutathione (hydro)per/polysulfides should be tightly controlled in subcellular compartments containing active glutaredoxins and high GSH concentrations.

## Introduction

Polysulfides (RS_n≥3_Rꞌ), hydropolysulfides (RS_n≥3_H) and hydropersulfides (RSSH) are versatile electron donors and acceptors with potential regulatory or cytoprotective physiological effects.^1–6^ For example, glutathione hydropersulfide (GSSH) not only has an increased nucleophilicity due to an α-effect but is also much more acidic than reduced glutathione (GSH), resulting in a higher availability of the nucleophilic anion for S_N_2 reactions.^7^ Furthermore, in contrast to thiols, stepwise oxidation of hydropersulfides results in reducible sulfur species, such as perthiosulfinic (RSSO_2_H) or perthiosulfonic acids (RSSO_3_H).^8, 9^ The outer sulfur atom of hydropersulfides could therefore serve as a dispensable protective cap that can be removed after alkylation or oxidative challenge.^9–11^ Hydropersulfides are also much better single-electron donors than the corresponding thiols and can act as physiological scavengers to disrupt radical chain reactions and protect membranes from lipid peroxidation.^4, 5, 12^ For example, GSSH was shown to react with lipid radicals in a radical-to-persulfide cycle (Fig. 1A).^5^ In this cycle, two glutathione perthiyl radicals (GSS^●^) can form glutathione tetrasulfide (GS_4_G), which reacts *in vitro* with 2 GSH, yielding GSSG and 2 GSSH. However, it is currently unclear how GS_4_G and related polysulfides are exactly reduced by GSH and whether enzymes could play a central role in these processes.^12^ Furthermore, glutathione-dependent enzymes could also affect the steady-state concentration of GSSH and other hydropersulfides, which were reported to reach concentrations greater than 0.1 mM.^1, 9^

**Figure 1.**
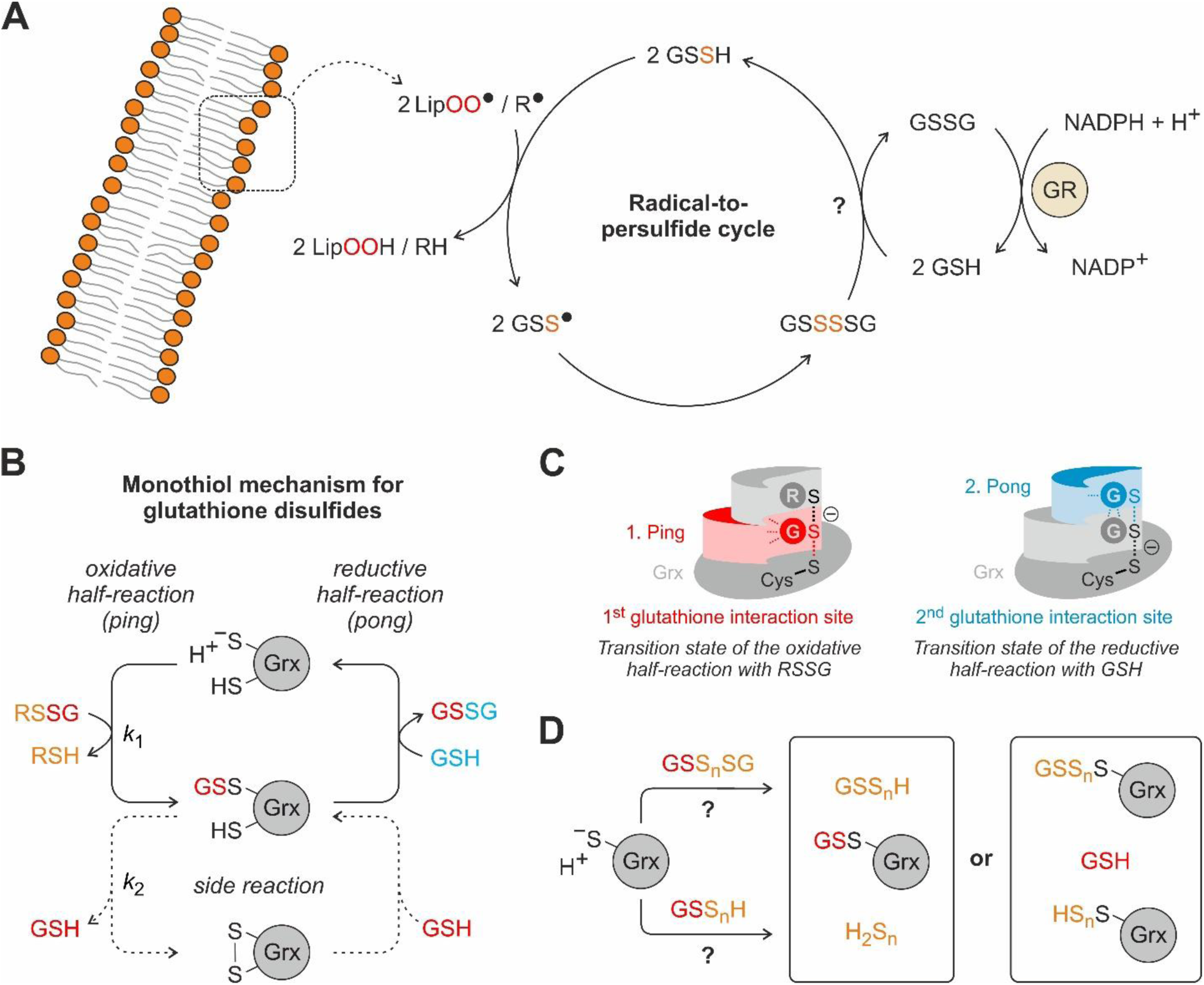
Potential relevance and mechanisms for the glutaredoxin-dependent reduction of glutathione di-, hydroper- and polysulfides. **A**) Membrane-protective radical-to-persulfide cycle as a sink for GSSH and source for GS_4_G. The rate constants and exact reduction sequence remain to be determined. GR, glutathione reductase; LipO^●^, lipid radical. **B**) Established reaction sequence for the reduction of mixed glutathione disulfides (RSSG) via a monothiol ping-pong mechanism, which requires the active-site cysteinyl thiolate (top). Some dithiol class I glutaredoxins readily form an intramolecular disulfide as a side reaction for RSSG substrates (bottom). **C**) Schematic representation of the predicted transition states of the monothiol ping-pong mechanism utilizing two different glutathione interaction sites for RSSG and GSH during the oxidative and reductive half-reaction, respectively. **D**) Theoretical mechanisms for the reduction of glutathione polysulfides (top) or hydroper/polysulfides (bottom). Depending on which sulfur atom is attacked by the active-site thiolate, the reactions can potentially yield either the glutathionylated enzyme (middle) or GSH (right).

Class I glutaredoxins are thiol:disulfide oxidoreductases that efficiently reduce a plethora of protein and low-molecular-weight disulfides.^13–23^ Common substrates are mixed glutathione disulfides (RSSG),^15, 17, 24, 25^ which are recognized at the so-called 1^st^ glutathione interaction site of the enzyme (Fig. 1B,C).^21, 22, 26–28^ The active-site cysteinyl thiolate serves as a nucleophile and attacks the *S*-glutathionylated substrate during the oxidative half-reaction, yielding a thiol product and the glutathionylated enzyme.^17, 21, 27, 28^ Deprotonated GSH then attacks the outer sulfur atom of the glutathionylated enzyme during the reductive half-reaction, yielding glutathione disulfide (GSSG) and the enzyme thiolate as an excellent leaving group (Fig. 1B,C).^17, 21, 25, 27, 29^ In contrast to the glutathione moiety of RSSG, GSH is recruited by an unspecific 2^nd^ glutathione interaction site (Fig. 1C).^21, 26, 28, 30^ Although all of the reactions in Fig. 1B are in principle reversible and glutaredoxins react with many thiols,^16, 29, 30^ the high intracellular concentration of GSH and the constant removal of GSSG by glutathione reductase usually result in a physiological GSH:GSSG ratio ≥10000:1,^31, 32^ which shifts the steady-state equilibrium towards the products and prevents the accumulation of glutathionylated proteins.^33^ Here, we used stopped-flow kinetic measurements in combination with mass spectrometry to address whether and how glutaredoxins also react with glutathione polysulfides (GS_n≥3_G), glutathione hydropolysulfides (GS_n≥3_H) and GSSH, summarized from hereon as glutathione (hydro)per/polysulfides, with implications for the steady-state concentrations and (potential) roles of these versatile redox metabolites (Fig. 1D).

## Results

### Mono- and dithiol class I glutaredoxins rapidly reduce GSSG

First, we directly determined the rate constant for the glutaredoxin-dependent reduction of GSSG as a reference. We therefore used previously established stopped-flow measurements for recombinant tryptophan mutants of the dithiol *P. falciparum* enzyme PfGrx^E28W/C88S^ (PfGrx^DT^), which still has both active-site cysteinyl residues C29 and C32, and its monothiol mutant PfGrx^E28W/C32S/C88S^ (PfGrx^MT^), which only has the catalytic cysteinyl residue C29.^30, 34^ The introduced tryptophan fluorophore is present at the same position in several other class I glutaredoxins^35, 36^ and was shown to have no or only little effect on the second-order rate constants for low-molecular-weight substrates tested so far.^30, 34^ In accordance with previous measurements,^34^ mixing reduced PfGrx^MT^ or PfGrx^DT^ with variable concentrations of GSSG resulted in a fast, concentration-dependent decrease in fluorescence during the first 100 milliseconds (Fig. 2A, Suppl. Fig. S1A). In contrast, the cysteine-free negative control PfGrx^E28W/C29S/C32S/C88S^ (PfGrx^NC^) revealed no change in fluorescence in the presence of GSSG (Suppl. Fig. S1B). Exponential fits of the kinetic traces yielded a second-order rate constant *k*_1_ of 4.1×10^6^ M^−1^s^−1^ for PfGrx^MT^ and 1.2×10^7^ M^−1^s^−1^ for PfGrx^DT^ (Fig. 2B, Table 1). Whole-protein mass spectrometry confirmed PfGrx^MT^(SSG) as the product of the reaction with GSSG (Fig. 2C). We therefore assign *k*_1_ to the glutathionylation of the active-site thiolate (Fig. 2D), which is in good agreement with rate constants obtained for the GSSG-dependent oxidation of roGFP2-fusion constructs of PfGrx without the tryptophan fluorophore at position 28 (Table 1).^22^ For PfGrx^DT^, we detected a second, much slower decrease in fluorescence for several seconds (right panel Fig. 2A). Rate constant *k*_2_ of this phase, which was around 0.28 s^−1^, did not depend on the GSSG concentration (right panel Fig. 2B, Table 1). We therefore assign the second phase to the formation of the intramolecular disulfide bond in PfGrx^DT^(S_2_) (Fig. 1B, see also below). For PfGrx^MT^, we observed a second, GSSG-dependent phase between 0.1–1 seconds with a very small change in fluorescence and a rate constant that might reflect the reverse reaction (left panel Fig. 2A, Suppl. Fig. S1C). To determine the relevance of the glutathione moiety of the disulfide substrate, we also analyzed the reduction of cystine by PfGrx^MT^, which revealed a small change in tryptophan fluorescence with a rate constant of ∼64 M^−1^s^−1^ in accordance with a non-enzymatic thiol-disulfide exchange reaction (Suppl. Fig. S2A,B, Table 1).^33^ In summary, the presence of the second cysteinyl residue at position 32 is not essential but increases the reactivity of the PfGrx active-site thiolate with GSSG, resulting in a reference rate constant *k*_1_ for the glutathionylation of PfGrx^DT^ on the order of 10^7^ M^−1^s^−1^.

**Figure 2.**
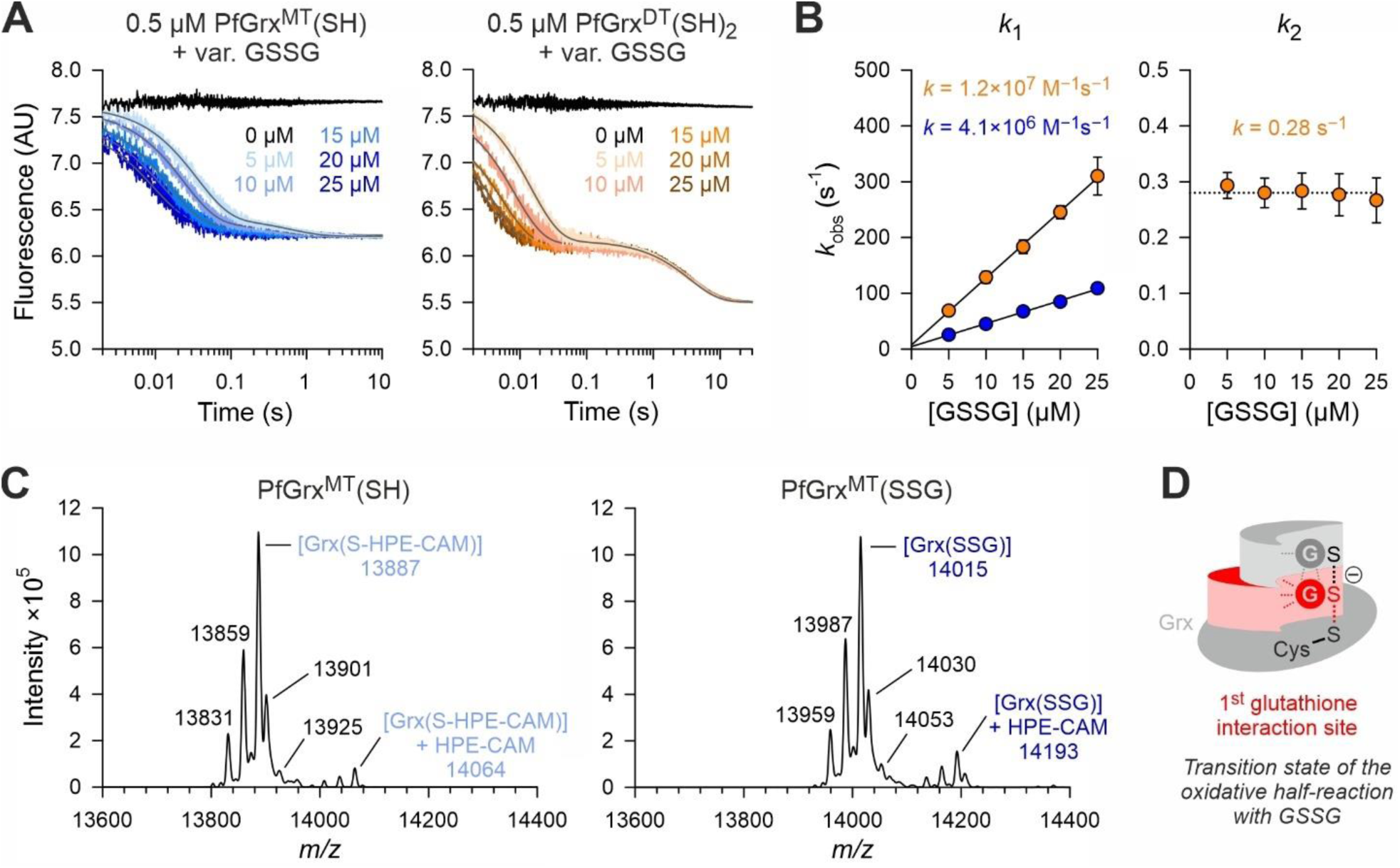
Direct rapid reduction of GSSG by PfGrx. **A**) Representative stopped-flow kinetics for the reaction of reduced PfGrx^MT^ (left) or PfGrx^DT^ (right) with variable concentrations of GSSG at 25°C and pH 7.4. **B**) Secondary plots for the observed rate constants (*k*_obs_) from double-exponential fits from panel A. The GSSG-dependent rate constant for the glutathionylation during the first phase (left) was determined from the slopes of the secondary plot, whereas the GSSG-independent rate constant for the intramolecular PfGrx^DT^ disulfide formation during the second phase (right) was averaged from all data points. Representative traces in panel A were averaged from three technical replicates and *k*_obs_ values in panel B were generated from three independent biological replicates (*n* = 3×3). **C**) Whole-protein mass spectrometry for reduced (left) and GSSG-oxidized (right) PfGrx^MT^ with calculated masses of 13888 and 14016 Da, respectively (following alkylation of free thiol groups with 0.5 mM HPE-IAM). Additional peaks may arise from oxidation of argininyl to glutamyl residues (–27 Da), methylation (+14 Da), potassium adducts (+38) or alkylation of amines by HPE-IAM (+178 Da). **D**) Schematic representation of the predicted transition state for the reaction between GSSG and the reduced enzyme.

**Table 1.** Rate constants for selected reactions of glutaredoxins and glutathione reductases.

| Assigned reaction partners A/B and products P/Q | | | | $y_0^a$ | Rate constant | pH/T |
| --- | --- | --- | --- | --- | --- | --- |
| A | B | P | Q |  |  |  |
| PfGrx<sup>MT</sup>(SH) | GSSG | PfGrx<sup>MT</sup>(SSG) | GSH | 4.5 s<sup>-1</sup> | 4.1×10<sup>6</sup> M<sup>-1</sup>s<sup>-1</sup> | 7.4/25°C |
| roGFP2-PfGrx<sup>C32S/C88S</sup>(SH) | GSSG | roGFP2-PfGrx<sup>C32S/C88S</sup>(SSG) | GSH | - | 2.6×10<sup>6</sup> M<sup>-1</sup>s<sup>-1</sup> <sup>b</sup> | 8.0/25°C |
| PfGrx<sup>DT</sup>(SH)<sub>2</sub> | GSSG | PfGrx<sup>DT</sup>(SSG)(SH) | GSH | 7.5 s<sup>-1</sup> | 1.2×10<sup>7</sup> M<sup>-1</sup>s<sup>-1</sup> | 7.4/25°C |
| roGFP2-PfGrx<sup>C88S</sup>(SH)<sub>2</sub> | GSSG | roGFP2-PfGrx<sup>C88S</sup>(SSG)(SH) | GSH | - | 1.4×10<sup>7</sup> M<sup>-1</sup>s<sup>-1</sup> <sup>b</sup> | 8.0/25°C |
| PfGrx<sup>DT</sup>(SSG)(SH) | - | PfGrx<sup>DT</sup>(S)<sub>2</sub> | GSH | - | 0.26–0.28 s<sup>-1</sup> | 7.4/25°C |
| PfGrx<sup>MT</sup>(SH) | GS<sub>3</sub>G | PfGrx<sup>MT</sup>(SSG) | GSSH | -21 s<sup>-1</sup> | 1.7×10<sup>7</sup> M<sup>-1</sup>s<sup>-1</sup> | 7.4/25°C |
| PfGrx<sup>DT</sup>(SH)<sub>2</sub> | GS<sub>3</sub>G | PfGrx<sup>DT</sup>(SSG)(SH) | GSSH | -12 s<sup>-1</sup> | 2.1×10<sup>7</sup> M<sup>-1</sup>s<sup>-1</sup> | 7.4/25°C |
| PfGrx<sup>MT</sup>(SH) | GS<sub>4</sub>G | PfGrx<sup>MT</sup>(SSG) | GS<sub>3</sub>H | -4.5 s<sup>-1</sup> | ~2.2×10<sup>7</sup> M<sup>-1</sup>s<sup>-1</sup> | 7.4/25°C |
| PfGrx<sup>DT</sup>(SH)<sub>2</sub> | GS<sub>4</sub>G | PfGrx<sup>DT</sup>(SSG)(SH) | GS<sub>3</sub>H | -11 s<sup>-1</sup> | ~2.7×10<sup>7</sup> M<sup>-1</sup>s<sup>-1</sup> | 7.4/25°C |
| PfGrx<sup>MT</sup>(SH) | GSSH | PfGrx<sup>MT</sup>(SSG) | H<sub>2</sub>S | 9.0 s<sup>-1</sup> | 3.9×10<sup>6</sup> M<sup>-1</sup>s<sup>-1</sup> | 7.4/25°C |
| PfGrx<sup>DT</sup>(SH)<sub>2</sub> | GSSH | PfGrx<sup>DT</sup>(SSG)(SH) | H<sub>2</sub>S | 5.3 s<sup>-1</sup> | 7.2×10<sup>6</sup> M<sup>-1</sup>s<sup>-1</sup> | 7.4/25°C |
| PfGrx<sup>MT</sup>(SSG) | H<sub>2</sub>S | PfGrx<sup>MT</sup>(SH) | GSSH | 0.38 s<sup>-1</sup> | 6.4×10<sup>4</sup> M<sup>-1</sup>s<sup>-1</sup> | 7.4/25°C |
| PfGrx<sup>MT</sup>(SH) | GS<sub>3</sub>H <sup>c</sup> | PfGrx<sup>MT</sup>(SSG) | H<sub>2</sub>S<sub>2</sub> | - | ~10<sup>6</sup>–10<sup>7</sup> M<sup>-1</sup>s<sup>-1</sup> <sup>c</sup> | 7.4/25°C |
| PfGrx<sup>DT</sup>(SH)<sub>2</sub> | GS<sub>3</sub>H <sup>c</sup> | PfGrx<sup>DT</sup>(SSG)(SH) | H<sub>2</sub>S<sub>2</sub> | - | ~10<sup>6</sup>–10<sup>7</sup> M<sup>-1</sup>s<sup>-1</sup> <sup>c</sup> | 7.4/25°C |
| PfGrx<sup>MT</sup>(SH) | CysSSCys | PfGrx<sup>MT</sup>(SSCys) | Cys | 2.0×10<sup>-3</sup> s<sup>-1</sup> | ~64 M<sup>-1</sup>s<sup>-1</sup> | 7.4/25°C |
| PfGrx<sup>MT</sup>(SH) | CysS<sub>3</sub>Cys | PfGrx<sup>MT</sup>(SSCys) | CysSSH | 5.8×10<sup>-2</sup> s<sup>-1</sup> | 2.7×10<sup>3</sup> M<sup>-1</sup>s<sup>-1</sup> <sup>d</sup> | 7.4/25°C |
| PfGrx<sup>MT</sup>(SSG) | GSH | PfGrx<sup>MT</sup>(SH) | GSSG | 2.6 s<sup>-1</sup> | 2.5×10<sup>5</sup> M<sup>-1</sup>s<sup>-1</sup> <sup>e</sup> | 7.4/25°C |
| PfGrx<sup>C32S/C88S</sup>(SSG) | GSH | PfGrx<sup>C32S/C88S</sup>(SH) | GSSG | - | 1.2×10<sup>5</sup> M<sup>-1</sup>s<sup>-1</sup> <sup>f</sup> | 8.0/25°C |
| ScGrx7<sup>G107W</sup>(SSG) | GSH | ScGrx<sup>G107W</sup>(SH) | GSSG | 4.3 s<sup>-1</sup> | 2.6×10<sup>5</sup> M<sup>-1</sup>s<sup>-1</sup> <sup>e</sup> | 7.4/25°C |
| ScGrx7(SSG) | GSH | ScGrx(SH)<sub>2</sub> | GSSG | - | 3.1×10<sup>5</sup> M<sup>-1</sup>s<sup>-1</sup> <sup>f,g</sup> | 8.0/25°C |
| hGrx1(SSG)(SH) | GSH | hGrx1(SH)<sub>2</sub> | GSSG | - | 1.0×10<sup>5</sup> M<sup>-1</sup>s<sup>-1</sup> <sup>h</sup> | 7.5/30°C |
| hGrx2(SSG)(SH) | GSH | hGrx2(SH)<sub>2</sub> | GSSG | - | 7.2×10<sup>3</sup> M<sup>-1</sup>s<sup>-1</sup> <sup>h</sup> | 7.5/30°C |
| hGrx2<sup>C40S</sup>(SSG) | GSH | hGrx2<sup>C40S</sup>(SH) | GSSG | - | 1.6×10<sup>4</sup> M<sup>-1</sup>s<sup>-1</sup> <sup>h</sup> | 7.5/30°C |
| PtGrxS12(SSG) | GSH | PtGrxS12(SH) | GSSG | - | 2.4×10<sup>4</sup> M<sup>-1</sup>s<sup>-1</sup> <sup>i</sup> | 7.9/25°C |
| ScGR + NADPH + H<sup>+</sup> | GSSG | ScGR + NADP<sup>+</sup> | 2 GSH | - | 3.9×10<sup>6</sup> M<sup>-1</sup>s<sup>-1</sup> <sup>j</sup> | 7.6/25°C |
| BtGR + NADPH + H<sup>+</sup> | GSSG | BtGR + NADP<sup>+</sup> | 2 GSH | - | 3.0×10<sup>6</sup> M<sup>-1</sup>s<sup>-1</sup> <sup>k</sup> | 7.25/28°C |
| BtGR + NADPH + H<sup>+</sup> | GS<sub>3</sub>G | BtGR + NADP<sup>+</sup> | GSSH + GSH | - | 2.9×10<sup>6</sup> M<sup>-1</sup>s<sup>-1</sup> <sup>k</sup> | 7.25/28°C |

<sup>a</sup> Y-axis intercept of secondary plots. Values must be interpreted with care, in particular, for single measurements with GS<sub>4</sub>G, GS<sub>3</sub>H or cystine.
<sup>b</sup> Second-order rate constants from stopped-flow measurements.<sup>22</sup>
<sup>c</sup> Reaction mixture contained significant amounts of GSSH besides GS<sub>3</sub>H.
<sup>d</sup> Equilibrium appears to be on the educt side and no PfGrx<sup>MT</sup>(SSCys) was detected by mass spectrometry.
<sup>e</sup> Second-order rate constants from stopped-flow measurements.<sup>30</sup>
<sup>f</sup> Reciprocal Dalziel-coefficients from steady-state measurements.<sup>21</sup>
<sup>g</sup> Reciprocal Dalziel-coefficients from steady-state measurements.<sup>28</sup>
<sup>h</sup> $k_{cat}/K_m$ values from steady-state measurements.<sup>25</sup>
<sup>i</sup> $k_{cat}/K_m$ value from steady-state measurements.<sup>19</sup>
<sup>j</sup> $k_{cat}/K_m$ value from steady-state measurements.<sup>59</sup>
<sup>k</sup> $k_{cat}/K_m$ values from steady-state measurements.<sup>40</sup>

### PfGrx rapidly converts GS_3_G and GS_4_G to GSSH and GS_3_H

Next, we analyzed the reduction of the glutathione polysulfides GS_3_G and GS_4_G by PfGrx (Fig. 3, Suppl. Fig. S3). We therefore mixed PfGrx^MT^ or PfGrx^DT^ with variable concentrations of the glutathione polysulfides in the stopped-flow spectrofluorometer. Similar decreases of tryptophan fluorescence were observed for GS_3_G and GS_4_G as for GSSG (Fig. 2A, Fig. 3A, Suppl. Fig. S3A). In contrast, no change in fluorescence was observed for PfGrx^NC^ in the presence of GS_3_G (Suppl. Fig. S1B). Rate constant *k*_1_ of PfGrx^MT^ and PfGrx^DT^ for the concentration-dependent first phase increased for the polysulfides compared to GSSG (Fig. 2B, Fig. 3B, Suppl. Fig. S3B, Table 1). As with GSSG, the presence of the second cysteinyl residue accelerated the reactions with GS_3_G and GS_4_G. In contrast to *k*_1_, *k*_2_ for PfGrx^DT^ was concentration-independent and similar regardless of the oxidant, which is consistent with a conversion of PfGrx^DT^(SSG) to PfGrx^DT^(S_2_) (Fig. 3B, Suppl. Fig. S3B). In accordance with a common mechanism and absent parallel reactions, whole-protein mass spectrometry revealed the formation of PfGrx^MT^(SSG) and PfGrx^DT^(S_2_) for both polysulfides, whereas no PfGrx(S_n≥3_Rꞌ) or PfGrx(S_n≥2_H) species were detected (Fig. 3C, Suppl. Fig. S3C). Furthermore, quantitative low-molecular-weight mass spectrometry confirmed the formation of GSSH and GS_3_H from GS_3_G and GS_4_G, respectively (Fig. 3D, Suppl. Fig. S3D). (For PfGrx^DT^, the detected GSSH or GS_3_H concentrations were lower, presumably because PfGrx^DT^(S_2_) and GSH were formed: GSH subsequently reduced a fraction of the enzyme, which then reacted with GSSH or GS_3_H, yielding H_2_S and hydrogen polysulfides (H_2_S_n≥2_) even during the short incubation period as outlined below. GS_3_H could also react with GSH, yielding GSSG and H_2_S_2_, which could explain the formation of GSSH from GSH). We interpret the results in Fig. 3 and Suppl. Fig. S3 as a rapid enzyme-catalyzed reduction of GS_3_G or GS_4_G due to a specific recognition of the substrate glutathione moiety at the 1^st^ glutathione interaction site, yielding (irrespective of the oxidant) the glutathionylated enzyme and deprotonated GSSH or GS_3_H, respectively (Fig. 3E, Suppl. Fig. S3E). The latter are better leaving groups than deprotonated GSH, which explains the increase in *k*_1_. In contrast to GS_3_G, cysteine trisulfide (CysS_3_Cys) was reduced much more slowly by PfGrx^MT^ with a second order rate constant of 2.7×10^3^ M^−1^s^−1^ (and the fits required an additional phase with a second-order rate constant of 2.6×10^2^ M^−1^s^−1^) (Suppl. Fig. S2A,C, Table 1). Furthermore, as indicated by the smaller change in tryptophan fluorescence and the data from mass spectrometry, only little CysS_3_Cys was reduced and no cysteinylated PfGrx^MT^ was detected, suggesting that the reverse reaction with highly reactive cysteine hydropersulfide (CysSSH) is favored (Suppl. Fig. S2A,D,E). In summary, glutathione polysulfides are excellent glutaredoxin substrates and are reduced even faster than GSSG.

**Figure 3.**
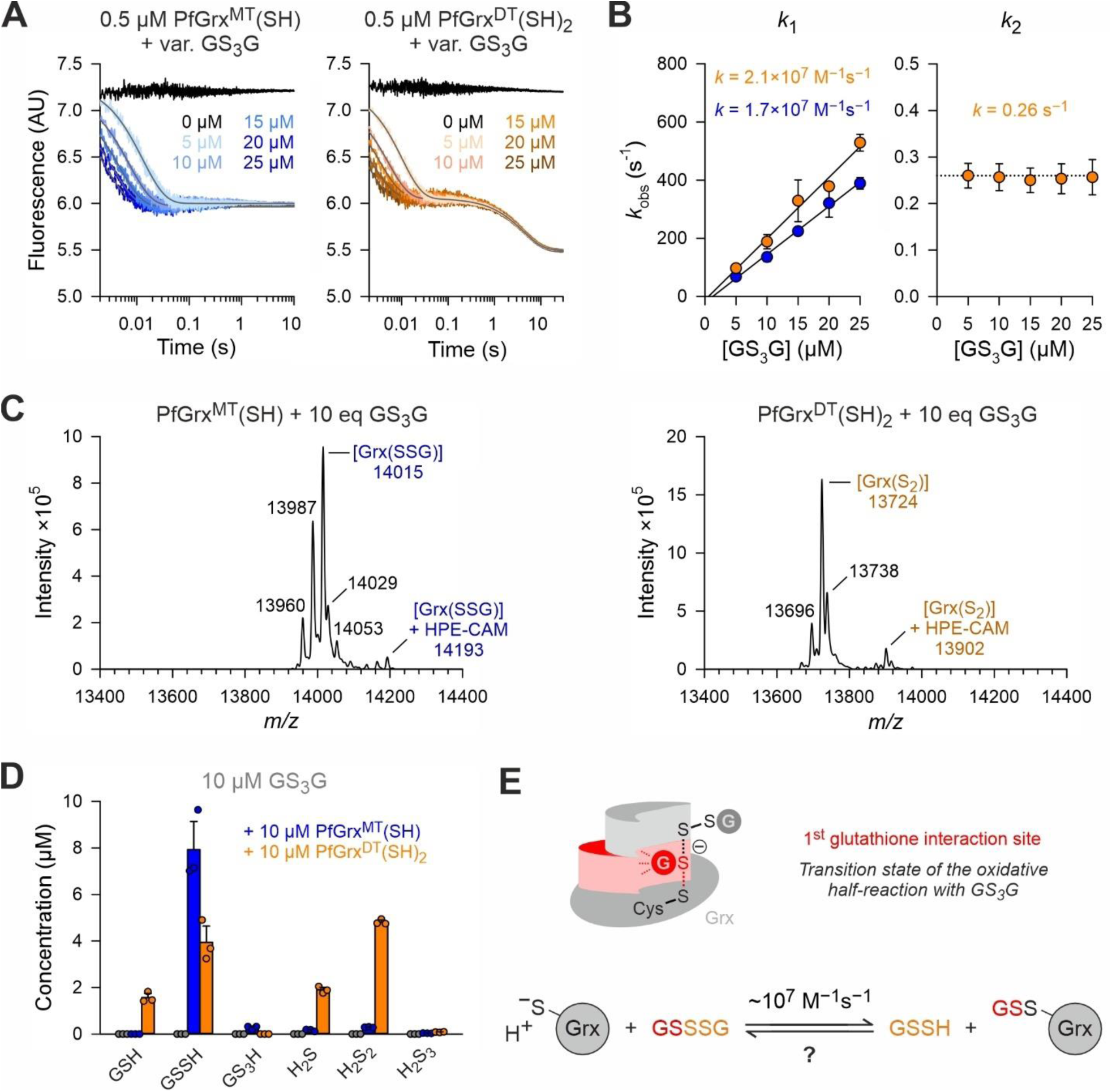
Direct rapid reduction of GS_3_G by PfGrx. **A**) Representative stopped-flow kinetics for the reaction of reduced PfGrx^MT^ (left) or PfGrx^DT^ (right) with variable concentrations of GS_3_G at 25°C and pH 7.4. **B**) Secondary plots for the *k*_obs_ values from single-or double-exponential fits from panel A. The GS_3_G-dependent rate constant for the first phase (left) was determined from the slopes of the secondary plot, whereas the GS_3_G-independent rate constant for the second phase for PfGrx^DT^ (right) was averaged from all data points. Representative traces in panel A were averaged from three technical replicates and *k*_obs_ values in panel B were generated from three independent biological replicates (*n* = 3×3). **C**) Whole-protein mass spectrometry after incubation of 10 equivalents (eq) of GS_3_G with reduced PfGrx^MT^ or PfGrx^DT^ for 10 s on ice. Calculated masses for glutathionylated PfGrx^MT^ or PfGrx^DT^(S_2_) are 14016 Da or 13725 Da, respectively (following treatment with 2 mM HPE-IAM). Additional peaks may arise from oxidation of argininyl to glutamyl residues (–27 Da), methylation (+14 Da), potassium adducts (+38) or alkylation of amines by HPE-IAM (+178 Da). **D**) Quantitative low-molecular-weight mass spectrometry after incubation of GS_3_G with either no enzyme, reduced PfGrx^MT^ or reduced PfGrx^DT^ for 10 s on ice. **E**) Schematic representation of the predicted transition state and assigned rate constant for the reaction between GS_3_G and the reduced enzyme.

### PfGrx rapidly converts GSSH and GS_3_H to H_2_S and H_2_S_2_

The results from Fig. 3 and Suppl. Fig. S3 revealed the glutaredoxin-dependent formation of GSSH and GS_3_H from glutathione polysulfides and already pointed towards a subsequent reduction of these products. To analyze the latter reaction in stopped-flow meas-urements, we generated GSSH and GS_3_H *in situ* using commercial yeast glutathione reductase (ScGR) with NADPH as electron donor (Fig. 4, Suppl. Fig. S4). Mixing ScGR and NADPH in the first syringe with GSSG, GS_3_G or GS_4_G in the second syringe revealed the consumption of one or up to two equivalents of NADPH (Fig. 4A, Suppl. Fig. S4A). The NADPH-oxidation kinetics were similar during the first three seconds, suggesting that ScGR forms GSH regardless of whether GSSG, GS_3_G or GS_4_G is the electron acceptor. A comparison of the kinetics in Fig. 3A, Suppl. Fig. S3A and Fig. 4A also suggests that recombinant PfGrx^DT^ reduces glutathione polysulfides faster than commercial ScGR. While GSSH and GS_3_H were formed in the presence of one equivalent of NADPH, mixing ScGR and two or three equivalents of NADPH in the first syringe with GS_3_G or GS_4_G in the second syringe resulted in an additional, much slower phase that was absent for GSSG (Fig. 4A, Suppl. Fig. S4A). Quantitative low-molecular-weight mass spectrometry supports our interpretation of the second phase as the ScGR-dependent reduction of the *in situ*-generated GSSH and GS_3_H, yielding a second molecule of GSH and either H_2_S or H_2_S_2_ (Fig. 4B,C Suppl. Fig. S4B,C). The data also confirms the instability of GS_3_H and formation of GSSH as outlined above. In summary, incubation of GS_3_G or GS_4_G with ScGR and one equivalent of NADPH results within seconds in the formation of GSSH or GS_3_H. ScGR can also reduce GSSH or GS_3_H at higher NADPH concentrations, however these reductions are much slower than with GS_3_G or GS_4_G.

**Figure 4.**
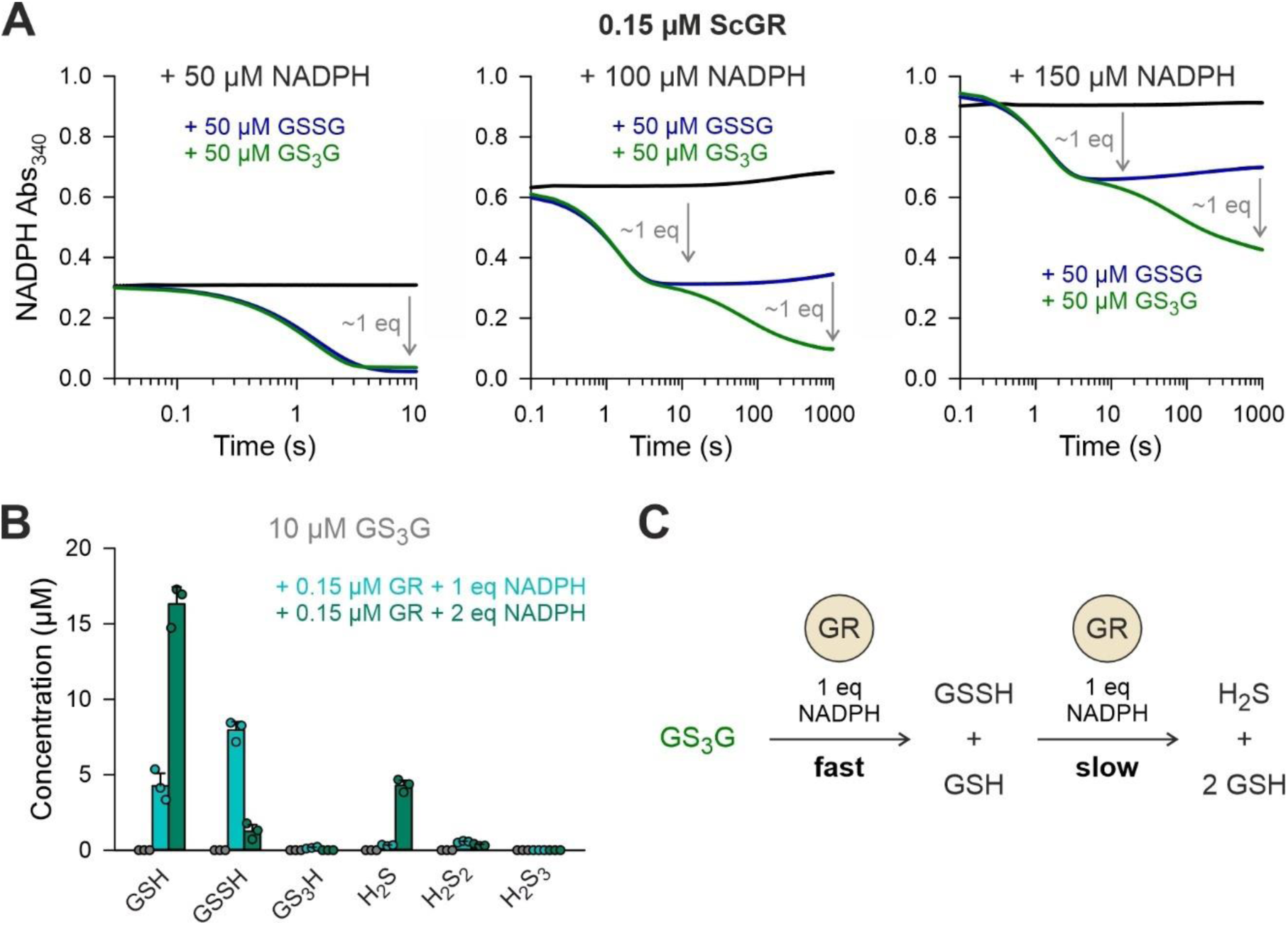
Glutathione reductase-dependent formation and reduction of GSSH. **A**) Representative stopped-flow kinetics for the reaction of ScGR with GSSG or GS_3_G in the presence of 1, 2 or 3 equivalents (eq) of NADPH at 25°C and pH 7.4. **B**) Quantitative low-molecular-weight mass spectrometry after incubation of GS_3_G with either no enzyme, ScGR and 1 eq NADPH (for 1 min on ice) or 2 eq NADPH (for 30 min at 25°C to allow the full reduction of GSSH). **C**) Schematic summary of the data interpretation.

Next, we shortly incubated GS_3_G or GS_4_G with ScGR and one equivalent of NADPH in the first syringe before mixing with reduced PfGrx^MT^ or PfGrx^DT^ in the second syringe (Fig. 5, Suppl. Fig. S5). We observed a decrease in tryptophan fluorescence in accordance with a glutathionylation of the enzyme by *in situ*-generated GSSH or GS_3_H (Fig. 5A, Suppl. Fig. S5A). In contrast, no change in fluorescence was observed for PfGrx^NC^ in the presence of GSSH (Suppl. Fig. S1B). Rate constant *k*_1_ for the GSSH- or GS_3_H-dependent first phase was 10^6^–10^7^ M^−1^s^−1^ for PfGrx^MT^ and PfGrx^DT^, which is up to one order of magnitude smaller than for the corresponding polysulfides (Fig. 5B, Suppl. Fig. S5B, Table 1). The presence of the second cysteinyl residue in PfGrx^DT^ again accelerated the reactions with GSSH and GS_3_H. (However, the rate constants for the reduction of GS_3_H must be interpreted with caution, as high amounts of GSSH and GSH were formed after the *in situ*-preparation of the substrate (Suppl. Fig. S5B-F). This could also explain why *k*_2_ for PfGrx^DT^ with GS_3_H was concentration-dependent, whereas *k*_3_ reflects the formation of PfGrx^DT^(S_2_) from PfGrx^DT^(SSG) (Suppl. Fig. S5C,D). Furthermore, the smaller decrease of tryptophan fluorescence in Suppl. Fig. S5A might point towards a pronounced backward reaction.) Whole-protein mass spectrometry confirmed again the formation of PfGrx^MT^(SSG) and PfGrx^DT^(S_2_) for both GSSH and GS_3_H, whereas no PfGrx(S_n≥3_Rꞌ) or PfGrx(S_n≥2_H) species were detected (Fig. 5C, Suppl. Fig. S6C, Suppl. Fig. S5E). In addition, quantitative low-molecular-weight mass spectrometry confirmed the ScGR-dependent formation and PfGrx-dependent consumption of GSSH and GS_3_H, yielding H_2_S and H_2_S_2_ (Fig. 5D, Suppl. Fig. S5F). Accordingly, we always detected the smell of H_2_S in samples when glutathione polysulfides were incubated for extended periods with NADPH and ScGR or active PfGrx variants. Since similar amounts of GSSH and GS_3_H were consumed in the presence of PfGrx in Suppl. Fig. S5F, both reactions probably proceeded with similar rate constants of 10^6^–10^7^ M^−1^s^−1^ (Table 1). We interpret the results in Fig. 5 and Suppl. Fig. S5 as a rapid enzyme-catalyzed reduction of GSSH and GS_3_H due to a specific recognition of the substrate glutathione moiety at the 1^st^ glutathione interaction site (Fig. 5E, Suppl. Fig. S5G). We also analyzed the reverse reaction for glutathionylated PfGrx^MT^ with variable concentrations of Na_2_S as a source for HS^−^. Secondary plots from single exponential fits of the increasing tryptophan fluorescence yielded a second-order rate constant of 6.4×10^4^ M^−1^s^−1^. We interpret this as a rather efficient reduction of PfGrx^MT^(SSG) by HS^−^, yielding the reduced enzyme and GSSH, as confirmed by whole-protein and low-molec-ular-weight mass spectrometry (Fig. 5F-H, Suppl. Fig. S6D). While the reduction of PfGrx^MT^(SSG) by HS^−^ was efficient but unspecific, PfGrx^DT^(S)_2_ was not efficiently reduced by HS^−^ (Suppl. Fig. S6D-F). These results can be explained by the two different substrates interaction sites in accordance with previous results for other thiols.^30^ In summary, GSSH and glutathione hydropolysulfides are more slowly reduced by glutathione reductase but are very good glutaredoxin substrates that are rapidly and reversibly converted to GSSG and H_2_S or the corresponding hydrogen polysulfides.

**Figure 5.**
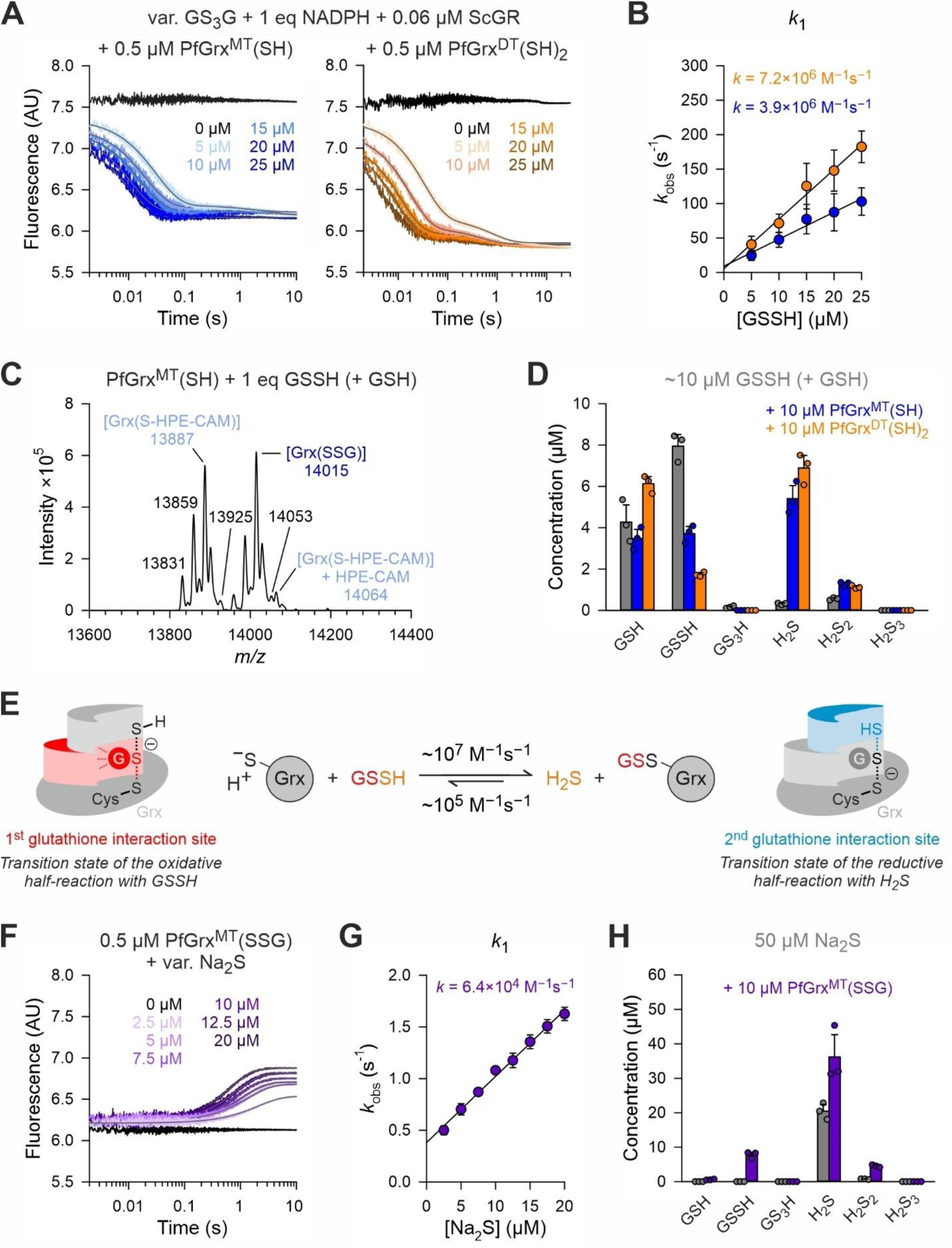
Direct rapid reduction of GSSH by PfGrx. **A**) Representative stopped-flow kinetics for the reaction of reduced PfGrx^MT^ (left) or PfGrx^DT^ (right) with variable concentrations of *in situ*-generated GSSH at 25°C and pH 7.4. **B**) Secondary plots for the *k*_obs_ values from double-exponential fits from panel A. The GSSH-dependent rate constant for the first phase was determined from the slopes of the secondary plot. Representative traces in panel A were averaged from three technical replicates and *k*_obs_ values in panel B were generated from three independent biological replicates (*n* = 3×3). **C**) Whole-protein mass spectrometry after incubation of *in situ*-generated GSSH with reduced PfGrx^MT^ for 10 s on ice. Calculated masses for reduced or glutathionylated PfGrx^MT^ are 13888 and 14016 Da, respectively (following treatment with 0.5 mM HPE-IAM). Additional peaks may arise from oxidation of argininyl to glutamyl residues (–27 Da), methylation (+14 Da), potassium adducts (+38) or alkylation of amines by HPE-IAM (+178 Da). **D**) Quantitative low-molecular-weight mass spectrometry after incubation of *in situ*-generated GSSH with either no enzyme, reduced PfGrx^MT^ or reduced PfGrx^DT^ for 10 s on ice. **E**) Schematic representation of the predicted transition state and assigned rate constants for the reversible reaction between GSSH and the reduced enzyme. **F**) Representative stopped-flow kinetics for the reaction of glutathionylated PfGrx^MT^ with variable concentrations of Na_2_S at 25°C and pH 7.4. **G**) Secondary plot for the *k*_obs_ values from single exponential fits from panel F. The Na_2_S-dependent rate constant was determined from the slope of the plot. Representative traces in panel F were averaged from three technical replicates and *k*_obs_ values in panel B were generated from three independent biological replicates (*n* = 3×3). **H**) Quantitative low-molecular-weight mass spectrometry after incubation of Na_2_S with either no enzyme or *S*-glutathionylated PfGrx^MT^ for 30 s on ice.

## Discussion

The steady-state concentrations of (hydro)per/polysulfides depend on their formation and conversion kinetics. This includes competing non-enzymatic as well as enzyme-catalyzed reactions. In order to better understand the physiological relevance of (hydro)per/polysulfides, we need to identify and characterize the most relevant reactions with the highest reaction rates at physiological metabolite concentrations.

What are the origins of GSSH and could glutaredoxins play a role in GSSH formation? The non-enzymatic formation of GSSH and GSH from H_2_S and GSSG has a rate constant of 0.2 M^−1^s^−^^1,7^ which is much too small to compete with the GSSG-dependent glutathionylation and H_2_S-dependent deglutathionylation of PfGrx^MT^ with rate constants of 4.1×10^6^ M^−1^s^−1^ and 6.4×10^4^ M^−1^s^−1^, respectively (Fig. 2A,B, Fig. 5F-H). However, even the glutaredoxin-catalyzed formation of GSSH and GSH from H_2_S and GSSG cannot be relevant under physiological conditions because GSH is usually the most abundant low-molecular-weight thiol in eukaryotes and many prokaryotes,^33, 37^ whereas H_2_S is usually in the nanomolar concentration range,^38, 39^ so that the steady-state equilibrium should be shifted towards the formation of GSSG instead of GSSH. In other words, a glutaredoxin-dependent formation of GSSH from GSSG would require toxic H_2_S concentrations and extremely small GSH:GSSG ratios. A far more likely source for GSSH and GS_n≥2_H appears to be the glutaredoxin- and/or glutathione reductase-dependent reduction of glutathione polysulfides (Fig. 3, Suppl. Fig. S3, Fig. 4, Suppl. Fig. S4),^10, 40^ although this necessitates polysulfides to start with. The radical-to-persulfide cycle in Fig. 1A provides one plausible scenario for the formation of glutathione polysulfides,^5^ but we currently neither know the exact concentrations nor rate constants of the relevant reactions. A non-enzymatic reduction of GS_4_G by GSH was also suggested but seems highly unlikely to be competitive considering an apparent rate constant of about 0.12 M^−1^s^−1^ for the reaction between GS_3_G and GSH.^41^ In the absence of polysulfides, GSSH can be generated by 3-mercaptopyruvate sulfur transferase, which forms an enzyme-bound hydropersulfide intermediate that can react with a variety of thiols including GSH.^42–44^ Furthermore, ubiquitous peroxiredoxin 6-type enzymes were suggested to convert H_2_O_2_ and H_2_S to H_2_S_2_, which might be another source for GSSH in eukaryotes and prokaryotes.^45^ Last but not least, hydropersulfides and H_2_S_2_ can be generated from prodrugs and could therefore also result in the formation of GSSH.^46, 47^ In summary, there are several, potentially competing (candidate) pathways for the formation of GSSH, including a rapid glutaredoxin-dependent reduction of glutathione polysulfides.

An enzyme system consisting of glutathione reductase, a glutaredoxin and GSH was previously shown to reduce polysulfides.^10^ Here we show that glutathione polysulfides and GS_n≥2_H are rapidly reduced by the model enzyme PfGrx with second-order rate constants ≥10^7^ M^−1^s^−1^ and 10^6^–10^7^ M^−1^s^−1^, respectively (Fig. 6, Table 1). All of these reactions proceed via a nucleophilic attack of the active-site thiolate at the sulfur atom of the glutathione moiety of the substrate, yielding only the glutathionylated enzyme and no other enzyme species. Glutaredoxins should therefore also efficiently reduce mixed polysulfides between glutathione and proteins. Glutathionylated glutaredoxins do not accumulate under physiological conditions because they are rapidly reduced by millimolar GSH and other thiols with second-order rate constants of 10^4^ –10^5^ M^−1^s^−1^ (Table 1).^19, 21, 25, 28–30^ Thus, glutaredoxins efficiently reduce glutathione polysulfides to GS_n≥2_H, which are subsequently converted to GSSG and H_2_S or the corresponding hydrogen polysulfides (Fig. 6). Glutathione reductase then reduces GSSG using NADPH as the electron donor, which shifts the steady-state equilibria towards the products. Bovine glutathione reductase was shown to also reduce GS_3_G with a *k*_cat_^app^/*K*_m_^app^ value of 2.9×10^6^ M^−1^s^−1^ at 28°C,^40^ which is about one order of magnitude slower than PfGrx^DT^ (Table 1). Thus, glutathione reductases might generally be less active than class I glutaredoxins. Considering also the abundances of both enzymes in various organisms and tissues,^33, 48^ we suggest that class I glutaredoxins often outcompete glutathione reductases for the reduction of glutathione polysulfides. In addition, as exemplified for PfGrx and ScGR, class I glutaredoxins clearly outcompete glutathione reductase for the reduction of GS_n≥2_H (Table 1) (although an enzymatic reduction might not be necessary *in vivo*, considering the apparent instability of GS_n≥3_H in the presence of millimolar GSH). As a consequence, glutathione (hydro)per/pol-ysulfides cannot accumulate under physiological conditions in the presence of active glutaredoxins and high GSH concentrations. How can we explain reported GSSH concentrations of 50 or even 150 µM, accounting for ≥1% of the total glutathione pool?^1^ On the one hand, subcellular compartments or microdomains lacking glutathione reductase and glutaredoxins^33^ might confound the GSSH value in total cell lysates. This scenario might somehow be comparable to initial reports on much too low GSH:GSSG ratios from invasive measurements in cell lysates, which were later revised with the help of genetically encoded redox sensors that allowed compartment-specific, non-invasive intracellular redox measurements.^31, 32^ On the other hand, high GSSH steady-state concentrations might originate from a very rapid and/or extensive formation of glutathione (hydro)per/polysulfides, e.g., in aerobic cell cultures with increased radical formation as shown in Fig. 1A or from a putative “storage pool” of non-glutathione (hydro)per/polysulfides. In contrast to GSSG and GS_3_G, cystine was not a substrate of PfGrx, while CysS_3_Cys was a moderate one, suggesting that glutaredoxins can, in principle, form CysSSH but that other proteins might be more relevant when cells are treated with CysS_3_Cys.^49–51^ Another interesting aspect of our results is that class I glutaredoxins should, in principle, increase the steady-state concentrations of H_2_S, H_2_S_2_ and other hydrogen polysulfides with implications for H_2_S-dependent redox signaling.^2, 38^ Glutaredoxins therefore antagonize the superoxide dismutase- or sulfide:quinone oxidoreductase-dependent decrease of the H_2_S steady-state concentration,^52–55^ as well as the non-enzymatic GSH-dependent consumption of H_2_S_2_ yielding GSSH (or consumption of H_2_S_n≥3_ yiedling the corresponding glutathione hydropolysulfides). In conclusion, class I glutaredoxins efficiently convert glutathione (hydro)per/polysulfides to GSSG and H_2_S or the corresponding hydrogen polysulfides.

**Figure 6.**
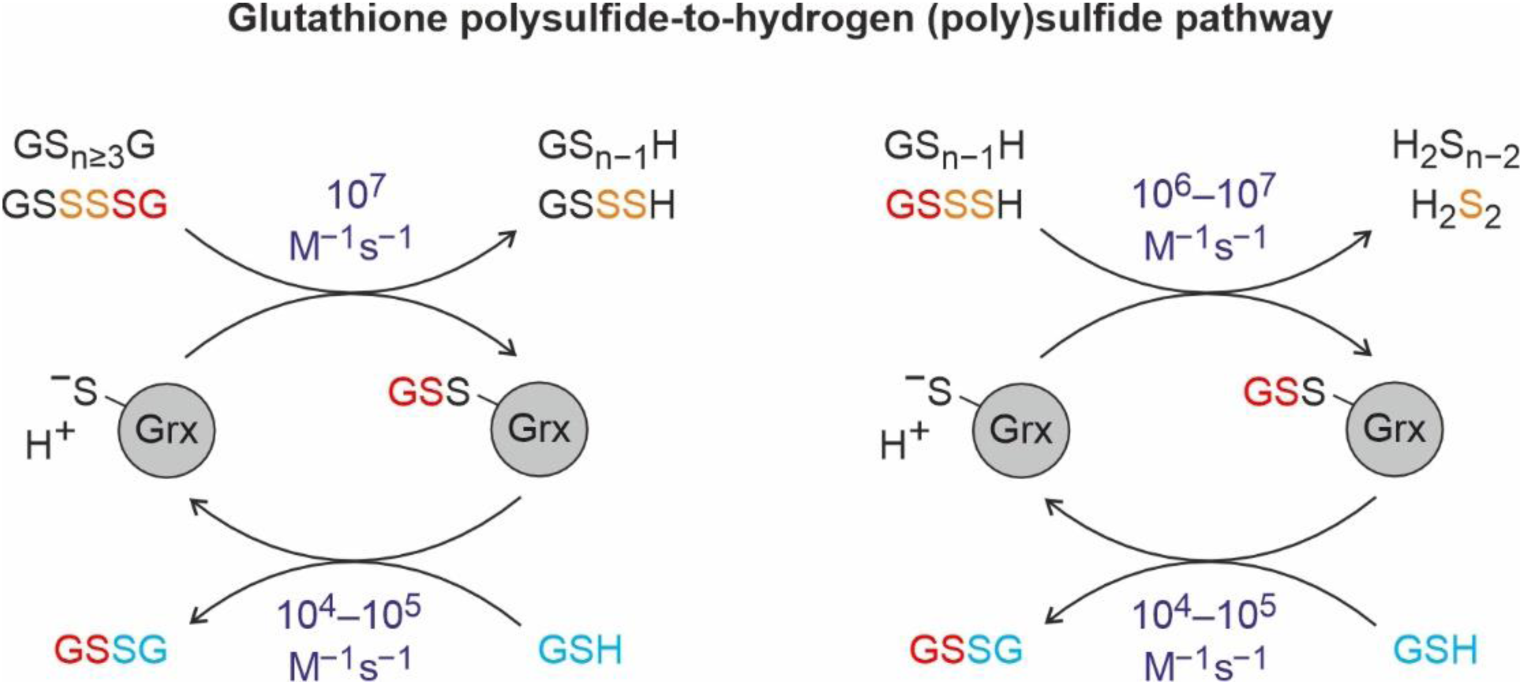
Glutaredoxin-catalyzed reduction of glutathione (hydro)per/polysulfides. Glutaredoxins could contribute to a membrane-protective radical-to-persulfide cycle and other persulfide pathways by rapidly reducing glutathione polysulfides to hydroper/polysulfides with second-order rate constants of ∼10^7^ M^−1^s^−1^. Glutathione hydroper/polysulfides could subsequently be reduced to H_2_S or hydrogen polysulfides by glutaredoxins with second-order rate constants of ∼10^6^–10^7^ M^−1^s^−1^. For example, GS_4_G is reduced by 2 GSH to 2 GSSG and H_2_S_2_.

## Materials and Methods

### Materials

The polysulfides GS_3_G, GS_4_G and CysS_3_Cys were synthesized as described previously.^1^ Diethylenetriaminepentaacetic acid (DTPA) was from Carl Roth, dithiothreitol (DTT), GSH, GSSG, 3-(dimethylcarbamoylimino)-1,1-dimethylurea (diamide), Na_2_S and ScGR were from Sigma-Aldrich, L-cystine was from Carbolution, NADPH was from Gerbu, *β*-(4-hydroxyphenyl)ethyl iodoacetamide (HPE-IAM) was from ChemCruz, and formic acid was from Merck. Primers were purchased from Metabion. All restriction enzymes, Phusion HF DNA polymerase, and T4 DNA ligase were from New England Biolabs.

### Site-directed mutagenesis

Plasmid pQE30/*PFGRX^C29S/C32S/C88S^* was generated by site-directed mutagenesis using *Pfu* polymerase (Promega) with pQE30/*PFGRX^C32S/C88S^* as a template,^56^ and mutagenesis primers C29S_forward (5′-GTATTTGCAAAAACGGAAAGCCCATATAGTATTAAGGC-3′) and C29S_reverse (5′-GCCTTAATACTATATGGGCTTTCCGTTTTTGCAAATAC-3′). The gene encoding PfGrx^NC^ was generated by site-directed mutagenesis using Phusion HF DNA polymerase with pQE30/*PFGRX^C29S/C32S/C88S^* as a template, and mutagenesis primers E28W_forward (5′-GCTGTATTTGCAAAAACGTGGAGCCCATATAGTATTAAGG-3′) and E28W_reverse (5′-CCTTAATACTATATGGGCTCCACGTTTTTGCAAATACAGC-3′). The methylated template DNA was digested with *Dpn*I, and the PCR product was transformed into chemically competent *E. coli* XL1-Blue cells. Plasmid DNA was isolated by minipreparation, and all constructs were verified by Sanger sequencing (Microsynth Seqlab).

### Heterologous expression, protein purification and sample preparation

Recombinant *N*-terminally MRGSH_6_GS-tagged PfGrx^DT^, PfGrx^MT^ and PfGrx^NC^ were produced in *E. coli* XL1-Blue cells and purified via affinity chromatography using nickel-nitrilotriacetic acid (Ni-NTA) agarose as described previously.^30, 34^ Fully reduced proteins were obtained by incubating the eluates from the Ni-NTA agarose column with 5 mM DTT for 30 min on ice. Excess DTT and imidazole were removed on a PD-10 desalting column (Cytiva) using 3.5 mL ice-cold assay-buffer (100 mM Na_x_H_y_PO_4_, 0.1 mM DTPA, pH 7.4 at 25 °C) for elution. To obtain PfGrx^DT^(S_2_) or PfGrx^MT^(SSG), the reduced proteins were incubated with 5 mM diamide or 5 mM GSSG for 1 h on ice and again purified using Ni-NTA agarose and PD-10 columns as described above. The purity of the final eluates was confirmed by analytical SDS-PAGE (Suppl. Fig. S7) and protein concentrations were determined spectrophotometrically at 280 nm. Extinction coefficients of 15.60 mM^−1^cm^−1^ for PfGrx^DT^(S)_2_ and 15.47 mM^−1^cm^−1^ for all other PfGrx variants were calculated from the amino-acid sequence using the Expasy ProtParam tool (https://web.expasy.org/prot-param).

### Fluorescence spectra

Fluorescence spectra of PfGrx^MT^ were recorded in a thermostatted SX-20 spectrofluorometer (Applied Photophysics) at 25°C using freshly purified enzyme. The tryptophan fluorescence was observed as the total emission at variable excitation wavelengths using a cut-off filter at 320 nm and a slit width of 2 mm. The spectra of reduced and glutathionylated PfGrx^MT^ were obtained 5 min after mixing 2 µM reduced PfGrx^MT^ in syringe 1 with either assay buffer or 120 µM GSSG in syringe 2.

### Stopped-flow fluorescence kinetic measurements

Stopped-flow fluorescence kinetic measurements were performed at 25°C using freshly purified enzyme. The change of tryptophan fluorescence was observed for up to 300 s as the total emission at an excitation wavelength of 295 nm using a cut-off filter at 320 nm and a slit width of 2 mm. The reaction of reduced PfGrx with GSSG, GS_3_G, GS_4_G, cystine or CysS_3_Cys was investigated by mixing 0.5–1 µM enzyme in syringe 1 with variable concentrations of substrate in assay buffer in syringe 2. Likewise, the reaction of reduced PfGrx with *in situ*-generated GSSH or GS_3_H was measured by mixing 0.5–1 µM enzyme in syringe 1 with 0.06 µM ScGR, variable concentrations of GS_3_G or GS_4_G and 1 equivalent NADPH in syringe 2. The incubation-time for the solution in syringe 2 prior to mixing was ∼1 min at 25°C to ensure the complete conversion of the polysulfides to hydroper/pol-ysulfides by ScGR. To investigate the reaction of oxidized PfGrx with Na_2_S, 1–2 µM PfGrx^DT^(S_2_) or PfGrx^MT^(SSG) in syringe 1 was mixed with variable concentrations of Na_2_S in syringe 2. Traces of 2–3 consecutive measurements were averaged and fitted by sin-gle-, double- or triple-exponential regression using the Pro-Data SX software (Applied Photophysics) to obtain *k*_obs_ values. The *k*_obs_ values of up to three biological replicates from independent protein purifications were averaged and plotted against the substrate concentration in SigmaPlot 13.0 to determine second-order rate constants from the slopes of the linear fits.

### Stopped-flow absorbance kinetic measurements

Stopped-flow absorbance kinetic measurements were performed at 25°C using ScGR. The consumption of NADPH was monitored at 340 nm for up to 1000 s using assay buffer as a blank, a slit width of 0.5 mm and an extinction coefficient of 6.22 mM^−1^cm^−1^. The reduction of GSSG, GS_3_G or GS_4_G by ScGR was investigated by mixing 0.1 mM substrate in syringe 1 with 1, 2 or 3 equivalents NADPH and 0.3 µM ScGR in syringe 2. Assays with assay buffer in syringe 1 served as negative controls. Traces of two consecutive measurements were averaged and plotted in SigmaPlot 13.0.

### Whole-protein mass spectrometry

Mass spectrometry samples were obtained following incubation of 10 µM reduced PfGrx^MT^ or PfGrx^DT^ with 10–250 µM substrate for up to 15 min on ice as indicated. GSSH or GS_3_H were prepared *in situ* as described above. Reference spectra were recorded for re-purified and desalted 10 µM reduced or *S*-glutathionylated PfGrx^MT^ as described above. All samples were incubated with 0.5 or 2 mM HPE-IAM for 20 min at 37°C. Samples were then diluted 1:10 with 0.1% formic acid and stored at –80°C. Whole-protein electrospray mass spectrometry was conducted in positive-ion mode using a liquid chromatography system with a POROS 10R1 column and a Bruker maXis time-of-flight spectrometer as described previously.^57^

### Quantitative low-molecular-weight mass spectrometry

Low-molecular-weight mass spectrometry samples for PfGrx were prepared as for whole-protein mass spectrometry. In case of ScGR, 0.15 µM enzyme was incubated with 10 µM GS_3_G or GS_4_G and 1 or 2 equivalents NADPH for up to 30 min at 25°C. Negative controls included samples without enzyme. All samples were incubated with 0.5 mM HPE-IAM for 20 min at 37 °C. Samples were then diluted 1:10 with 0.1% formic acid and stored at – 80°C. For quantitative low-molecular-weight mass spectrometry, samples were further diluted 1:1 with 0.1% formic acid containing known amounts of isotope-labeled internal standards. Analyses were performed on a LC-ESI-QTOF mass spectrometer (SYNAPT G2-Si, Waters) coupled to an ACQUITY UPLC system (Waters). Each sulfur metabolite was identified and quantified using targeted MS/MS mode. The synthesis of isotope-labelled internal standards, LC-ESI-MS conditions, and targeted MS/MS parameters were applied as described previously.^6, 58^

### Data availability

All relevant data are included in the paper or its Supporting information and are available from the authors upon reasonable request. The Supporting Information contains Figures S1 – S7. This material is available free of charge via the Internet at _xxx._

## Acknowledgements

The authors acknowledge Dr. Maria J. S. A. Silva, Dr. Seah Ling Kuan and the members of the MS core facility of the Max-Planck-Institute for Polymer Research for the support with mass spectrometry measurements and analysis in this project. We also thank Lukas Lang, Danny Schilling, Tobias Dick and Nicole Lübbehusen for suggestions and support with whole-protein mass spectrometry as well as Robin Schuhmann and Simon Unik for site-directed mutagenesis. The authors acknowledge the technical support of the Core Facility for Mass Spectrometry and Proteomics (CFMP) of ZMBH of Heidelberg. CFMP is funded by the ZMBH and partially funded by the CellNetworks Core Technology Platform (CCTP) of Heidelberg University. The CCTP is funded in part by the Federal Ministry of Education and Research (BMBF) and the Ministry of Science Baden Württemberg within the framework of the Excellence Strategy of the Federal and State Governments of Germany. This work was funded by the Deutsche Forschungsgemeinschaft (DFG) grant DE 1431/20-1 to M.D (project number 526346008) and, in part, RTG 2737 (project number 446816136). Open access funding enabled and organized by Projekt DEAL.

## Author Contributions

P.R. and L.L. performed all experiments and analyzed the data except for low-molecular-weight mass spectrometry which was performed and analyzed by S.O. and S.Y.. T.A. provided compounds and supervised the low-molecular-weight mass spectrometry analysis. U.B. and M.D. conceived and supervised the study. U.B. and M.D. wrote the manuscript. All authors discussed the results and gave approval to the final version of the manuscript.

