## Supplementary Data for "Glutaredoxins rapidly reduce glutathione hydroper- and polysulfides"

**Figure S1**

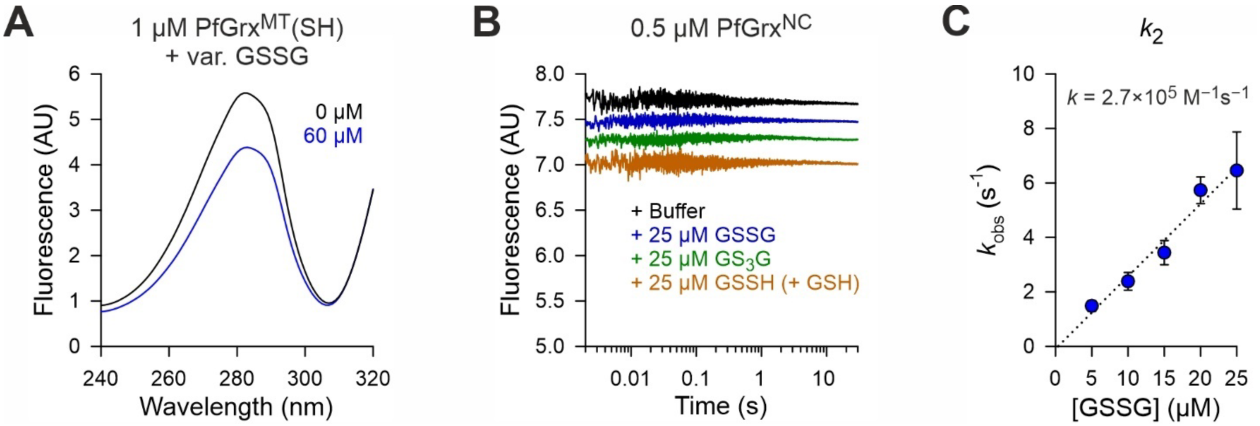


**Supplementary Figure S1 | Control experiments for stopped-flow measurements.** **A**) Representative fluorescence spectra of reduced and glutathionylated $PfGrx^{MT}$ recorded with the stopped-flow spectrofluorometer. **B**) Representative stopped-flow kinetics for the absent reaction of cysteine-free $PfGrx^{NC}$ with the indicated oxidants at 25°C and pH 7.4. **C**) Rate constant for a second GSSG-dependent reaction phase of $PfGrx^{MT}$ in Fig. 2A, which might reflect the reverse reaction.

### Figure S2

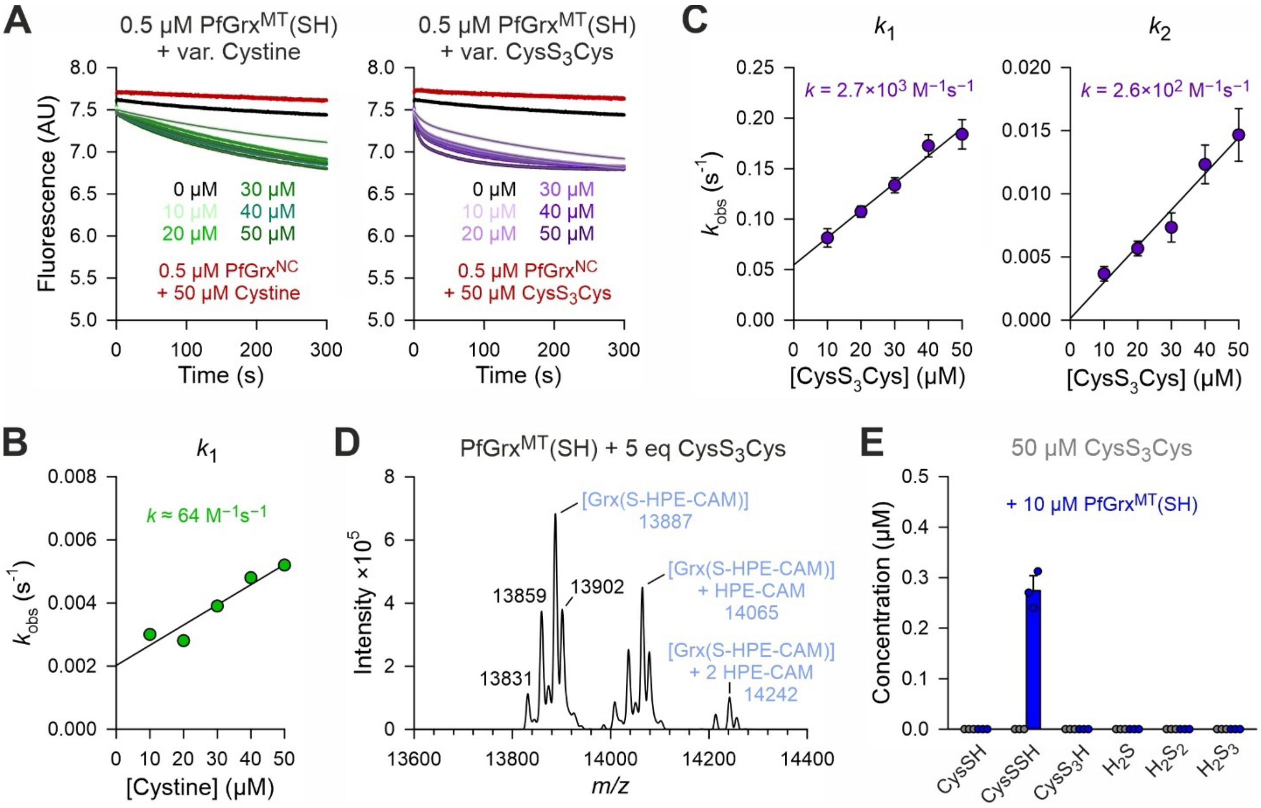


**Supplementary Figure S2 | Reaction kinetics for the $PfGrx^{MT}$-dependent reduction of cystine and $CysS_3Cys$. A**) Representative stopped-flow kinetics for the reaction of reduced $PfGrx^{MT}$ with variable concentrations of cystine (left) or $CysS_3Cys$ (right) at 25°C and pH 7.4. Control experiments with cysteine-free $PfGrx^{NC}$ are shown in red. **B**) Secondary plot for the $k_{obs}$ values from single exponential fits from panel A. The cystine-dependent rate constant was determined from the slope of the secondary plot. Representative traces in panel A were averaged from two technical replicates and $k_{obs}$ values in panel B were from a single experiment ($n$ = 1×2). **C**) Secondary plots for the $k_{obs}$ values from single exponential fits from panel A. The $CysS_3Cys$-dependent rate constant was determined from the slope of the secondary plots. Representative traces in panel A were averaged from two technical replicates and $k_{obs}$ values in panel C were generated from three independent biological replicates ($n$ = 3×2). **D**) Whole-protein mass spectrometry after incubation of $CysS_3Cys$ with reduced $PfGrx^{MT}$ for 10 s on ice. The calculated mass for reduced $PfGrx^{MT}$ is 13888 Da (following treatment with 2 mM HPE-IAM). Additional peaks may arise from oxidation of argininyl to glutamyl residues (–27 Da), methylation (+14 Da), potassium adducts (+38) or alkylation of amines by HPE-IAM (+178 Da). The peak at 13831, which is close to the calculated mass of cysteinylated $PfGrx^{MT}$ of 13830 Da, was also detected with a similar intensity for the reduced enzyme in Fig. 2C. **E**) Quantitative low-molecular-weight mass spectrometry after incubation of $CysS_3Cys$ with either no enzyme or reduced $PfGrx^{MT}$ for 15 min on ice.

### Figure S3

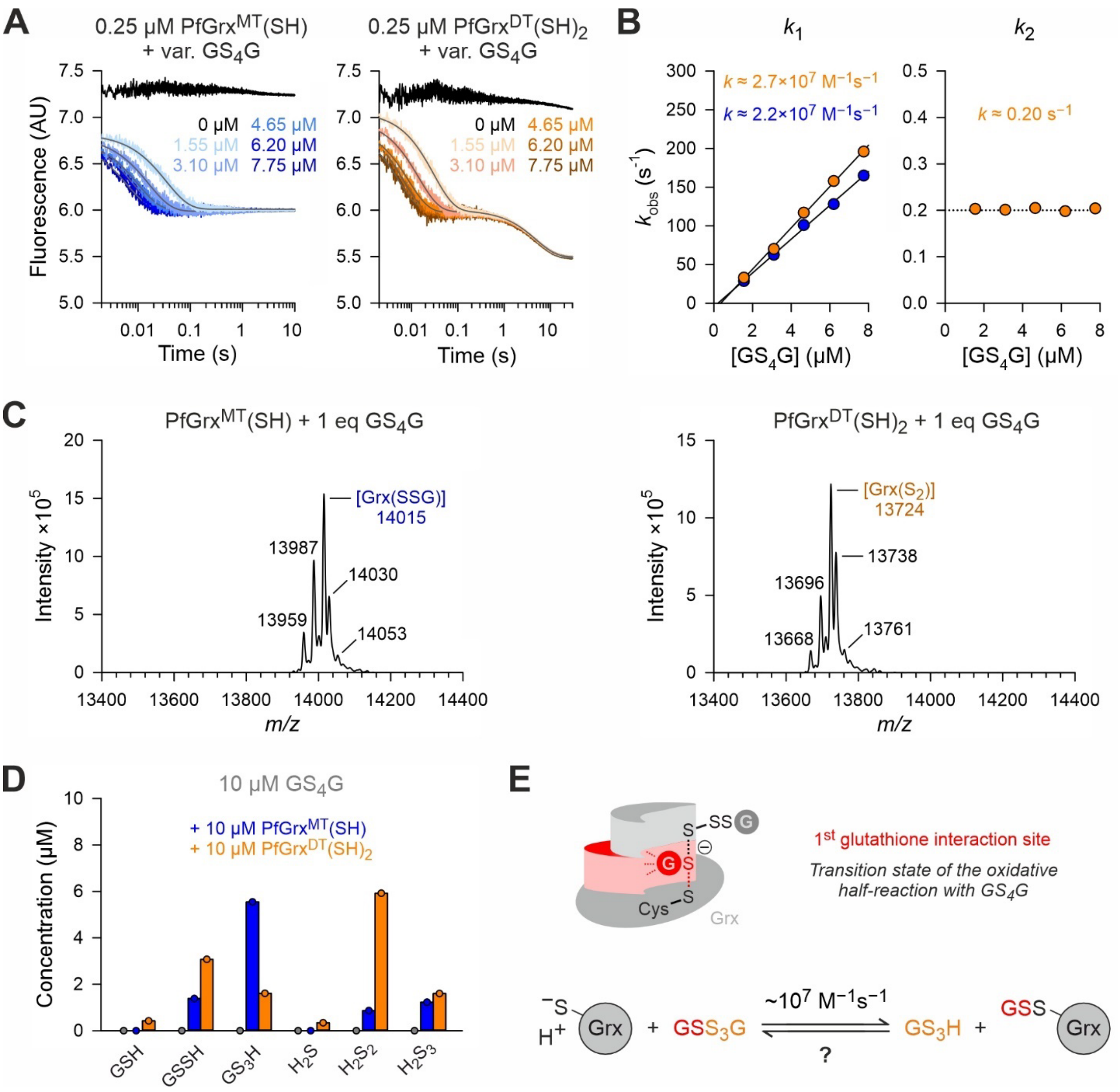


**Supplementary Figure S3 | Direct rapid reduction of $GS_4G$ by PfGrx. A**) Representative stopped-flow kinetics for the reaction of reduced $PfGrx^{MT}$ (left) or $PfGrx^{DT}$ (right) with variable concentrations of $GS_4G$ at 25°C and pH 7.4. **B**) Secondary plots for the $k_{obs}$ values from single- or double-exponential fits from panel A. The $GS_4G$-dependent rate constant for the first phase (left) was determined from the slopes of the secondary plot, whereas the $GS_4G$-independent rate constant for the second phase for $PfGrx^{DT}$ (right) was averaged from all data points. Representative traces in panel A were averaged from three technical replicates and $k_{obs}$ values in panel B were from a single experiment (*n* = 1×3). **C**) Whole-protein mass spectrometry after incubation of $GS_4G$ with reduced $PfGrx^{MT}$ or $PfGrx^{DT}$ for 10 s on ice. Calculated masses for glutathionylated $PfGrx^{MT}$ or $PfGrx^{DT}(S_2)$ are 14016 Da or 13725 Da, respectively (following treatment with 0.5 mM HPE-IAM). Additional peaks may arise from oxidation of argininyl to glutamyl residues (–27 Da), methylation (+14 Da), potassium adducts (+38) or alkylation of amines by HPE-IAM (+178 Da). **D**) Quantitative low-molecular-weight mass spectrometry after incubation of $GS_4G$ with either no enzyme, reduced PfGrxMT or reduced PfGrxDT for 10 s on ice. **E**) Schematic representation of the predicted transition state and assigned rate constant for the reaction between $GS_4G$ and the reduced enzyme.

**Figure S4**

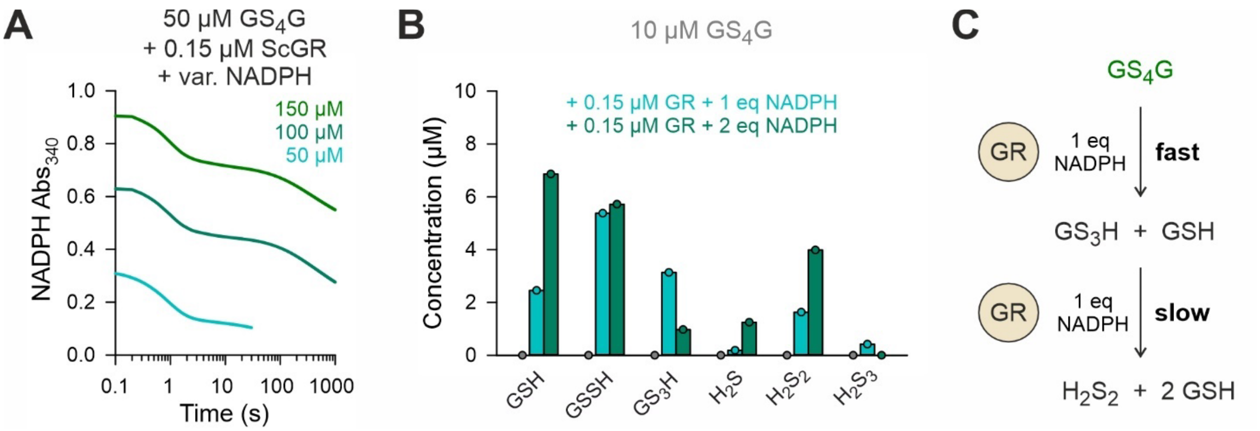


**Supplementary Figure S4 | Glutathione reductase-dependent formation and reduction of $GS_3H$. A**) Representative stopped-flow kinetics for the reaction of ScGR with $GS_4G$ in the presence of variable concentrations of NADPH at 25°C and pH 7.4. **B**) Quantitative low-molecular-weight mass spectrometry after incubation of $GS_4G$ with either no enzyme, ScGR and 1 eq NADPH (for 10 s on ice) or 2 eq NADPH (for 30 min at 25°C). **C**) Schematic summary of the data interpretation.

### Figure S5

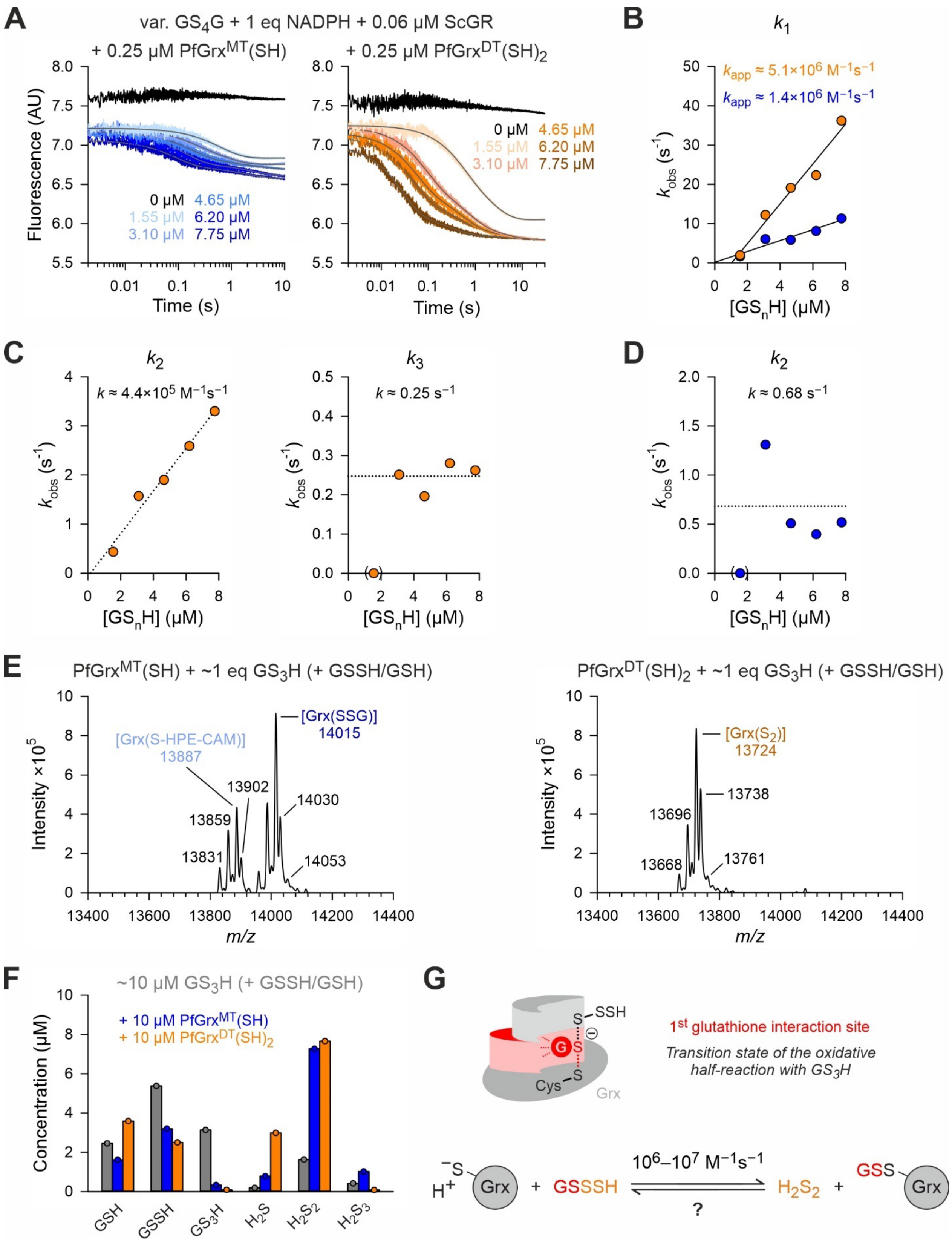

A
var. GS4G + 1 eq NADPH + 0.06 µM ScGR
+ 0.25 µM PfGrxMT(SH)
+ 0.25 µM PfGrxDT(SH)2
Fluorescence (AU)
Time (s)
0 µM 4.65 µM
1.55 µM 6.20 µM
3.10 µM 7.75 µM
B
k1
kapp ≈ 5.1×10^6 M−1s−1
kapp ≈ 1.4×10^6 M−1s−1
kobs (s-1)
[GSnH] (µM)
C
k2
k ≈ 4.4×10^5 M−1s−1
k3
k ≈ 0.25 s−1
D
k2
k ≈ 0.68 s−1
E
PfGrxMT(SH) + ~1 eq GS3H (+ GSSH/GSH)
PfGrxDT(SH)2 + ~1 eq GS3H (+ GSSH/GSH)
Intensity ×10^5
m/z
[Grx(S-HPE-CAM)] 13887
[Grx(SSG)] 14015
13902
13859
13831
14030
14053
[Grx(S2)] 13724
13738
13696
13668
13761
F
~10 µM GS3H (+ GSSH/GSH)
+ 10 µM PfGrxMT(SH)
+ 10 µM PfGrxDT(SH)2
Concentration (µM)
GSH
GSSH
GS3H
H2S
H2S2
H2S3
G
S SSH
S
Cys S
G
Grx
1st glutathione interaction site
Transition state of the oxidative half-reaction with GS3H
−S
H+
Grx
GSSSH
10^6–10^7 M−1s−1
?
H2S2
GSS
Grx

**Supplementary Figure S5 | Direct rapid reduction of $GS_3H$ by PfGrx. A**) Representative stopped-flow kinetics for the reaction of reduced $PfGrx^{MT}$ (left) or $PfGrx^{DT}$ (right) with variable concentrations of *in situ*-generated $GS_3H$ at 25°C and pH 7.4. **B–D**) Secondary plots for the $k_{obs}$ values from double- or triple-exponential fits from panel A. Actual concentrations may vary, as significant amounts of GSSH are formed besides $GS_3H$ during the pre-incubation of $GS_4G$ with GR and NADPH. The $GS_nH$-dependent rate constant was determined from the slopes of the secondary plot, whereas the $GS_nH$-independent rate constant for the third phase for $PfGrx^{DT}$ and the second phase for $PfGrx^{MT}$ was averaged from all data points. Representative traces in panel A were averaged from three technical replicates and $k_{obs}$ values in panel B–D were from a single experiment ($n$ = 1×3). **E**) Whole-protein mass spectrometry after incubation of *in situ*-generated $GS_3H$ with reduced $PfGrx^{MT}$ or $PfGrx^{DT}$ for 10 s on ice. Calculated masses for reduced and glutathionylated $PfGrx^{MT}$ or $PfGrx^{DT}(S_2)$ are 13888 Da and 14016 Da or 13725 Da, respectively (following treatment with 0.5 mM HPE-IAM). Additional peaks may arise from oxidation of argininyl to glutamyl residues (–27 Da), methylation (+14 Da), potassium adducts (+38) or alkylation of amines by HPE-IAM (+178 Da). **F**) Quantitative low-molecular-weight mass spectrometry after incubation of *in situ*-generated $GS_3H$ with either no enzyme, reduced $PfGrx^{MT}$ or reduced $PfGrx^{DT}$ for 10 s on ice. **G**) Schematic representation of the predicted transition state and assigned rate constant for the reaction between $GS_3H$ and the reduced enzyme.

### Figure S6

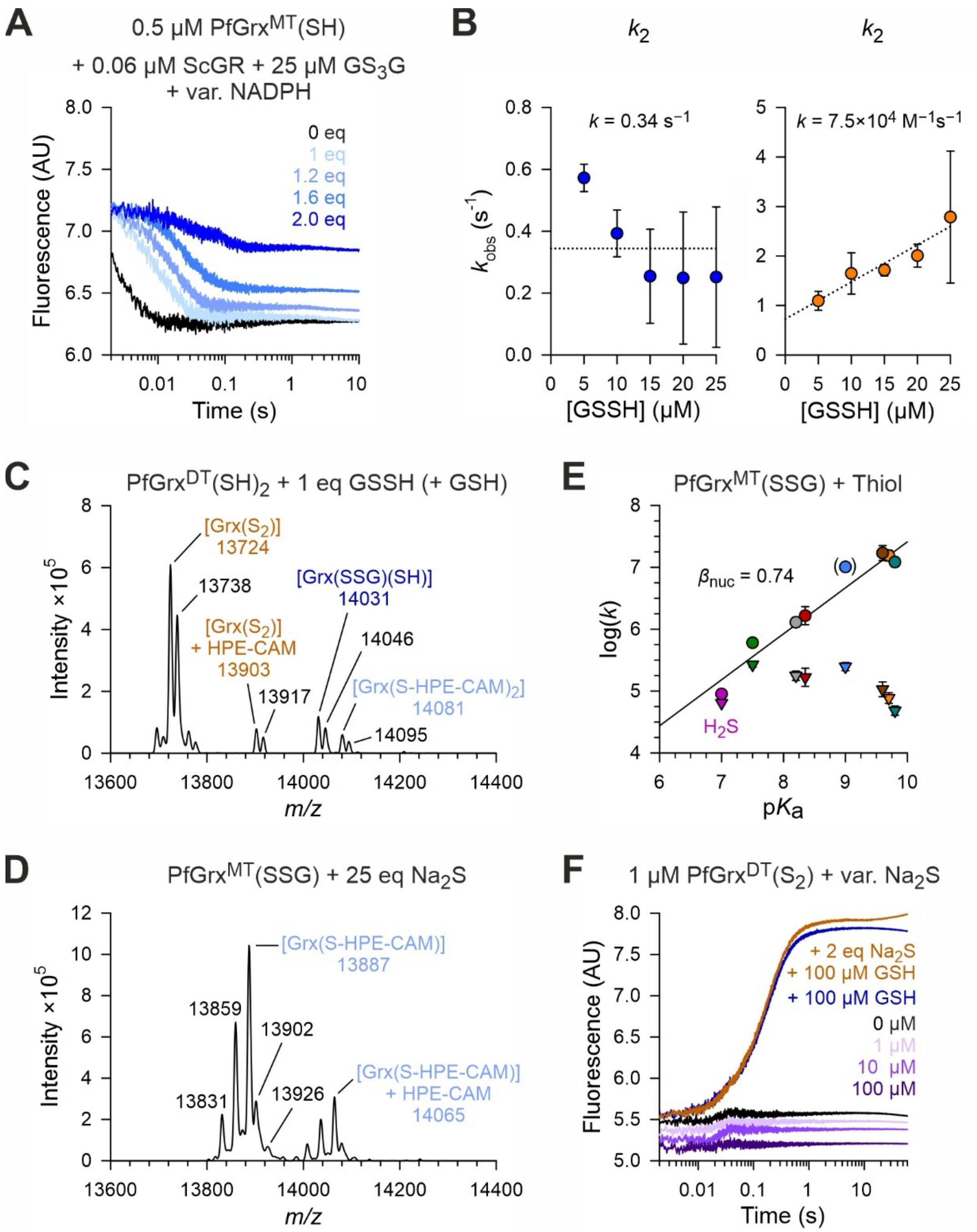


**Supplementary Figure S6 | Control experiments for the reduction of GSSH by PfGrx.** **A**) Representative stopped-flow kinetics for the reaction of reduced $PfGrx^{MT}$ with *in situ*-generated GSSH at 25°C and pH 7.4. Even after the addition of 2 eq NADPH, when all $GS_3G$ should be consumed, a fluorescence change was still observed, indicating that GSSH reacts directly with Grx. **B**) Rate constant for a second reaction phase of $PfGrx^{MT}$

(left) and $PfGrx^{DT}$ (right). **C**) Whole-protein mass spectrometry after incubation of *in situ*-generated GSSH with reduced $PfGrx^{DT}$ for 10 s on ice. Calculated masses for reduced and glutathionylated $PfGrx^{DT}$ or $PfGrx^{DT}(S)_2$ are 14081 Da and 14032 Da or 13725 Da, respectively (following treatment with 0.5 mM HPE-IAM). Additional peaks may arise from oxidation of argininyl to glutamyl residues (–27 Da), methylation (+14 Da), potassium adducts (+38) or alkylation of amines by HPE-IAM (+178 Da). **D**) Whole-protein mass spectrometry after incubation of $Na_2S$ with glutathionylated $PfGrx^{MT}$ for 15 min on ice. The calculated mass of reduced $PfGrx^{MT}$ is 13888 Da (following treatment with 2 mM HPE-IAM). **E**) Brønsted plot of the second-order rate constants (triangles) and normalized, pH-independent second-order rate constants (circles) for the reaction of $PfGrx^{MT}(SSG)$ with various thiols of differing p$k_a$-values determined previously.[1] The nucleophile Bronsted coefficient $\beta_{nuc}$ was obtained from the slope of the linear fit. The herein determined rate constant for $H_2S$ falls slightly below the linear regression, indicating no specificity of Grx(SSG) for $H_2S$ as a reducing agent compared to other thiols. **F**) Representative stopped-flow kinetics for the reaction of $PfGrx^{DT}(S)_2$ with various concentrations of $Na_2S$ (purple traces) at 25°C and pH 7.4. No fluorescence change was observed. The reduction by GSH served as a positive control (blue trace). Pre-incubation with $Na_2S$ did not significantly change the kinetics with GSH (orange trace), indicating that $Na_2S$ does not reduce $PfGrx^{DT}(S)_2$.

[1] Lang, L., Reinert, P., Diaz, C., and Deponte, M. (2024) The dithiol mechanism of class I glutaredoxins promotes specificity for glutathione as a reducing agent, *Redox Biol 78*, 103410.

**Figure S7**

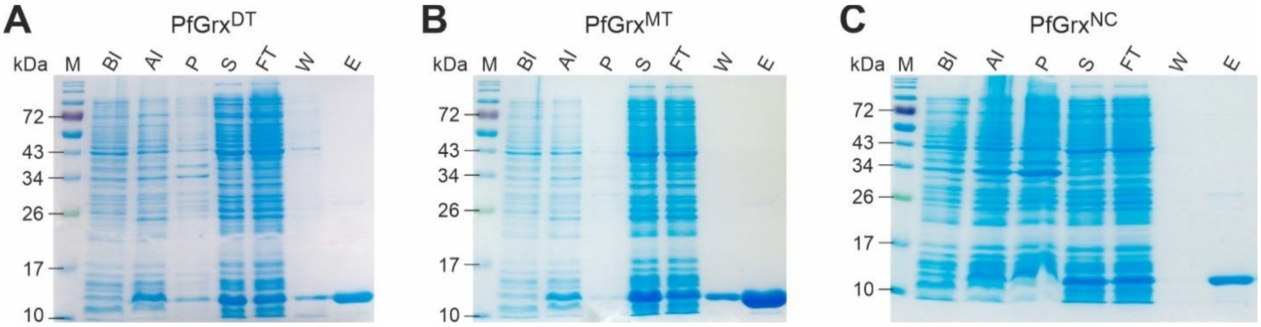


**Supplementary Figure S7 | SDS-PAGE analyses of representative purifications of recombinant PfGrx variants.** The indicated proteins were separated on a 15% SDS-PAGE gel under reducing conditions and visualized by Coomassie Brilliant Blue staining. Lanes: M, marker; BI, before induction; AI, after induction; P and S, pellet and supernatant after cell lysis and centrifugation; FT, Ni-NTA agarose flow through; W, Ni-NTA agarose wash fraction; E, Ni-NTA agarose eluate.
